## Supplementary figures and images for "ARL13B regulates Sonic Hedgehog signaling from outside primary cilia"

### Figure 2 Supplement 1

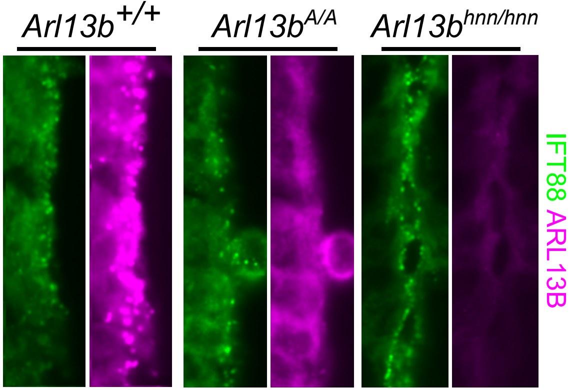

### Figure 3 Supplement 1

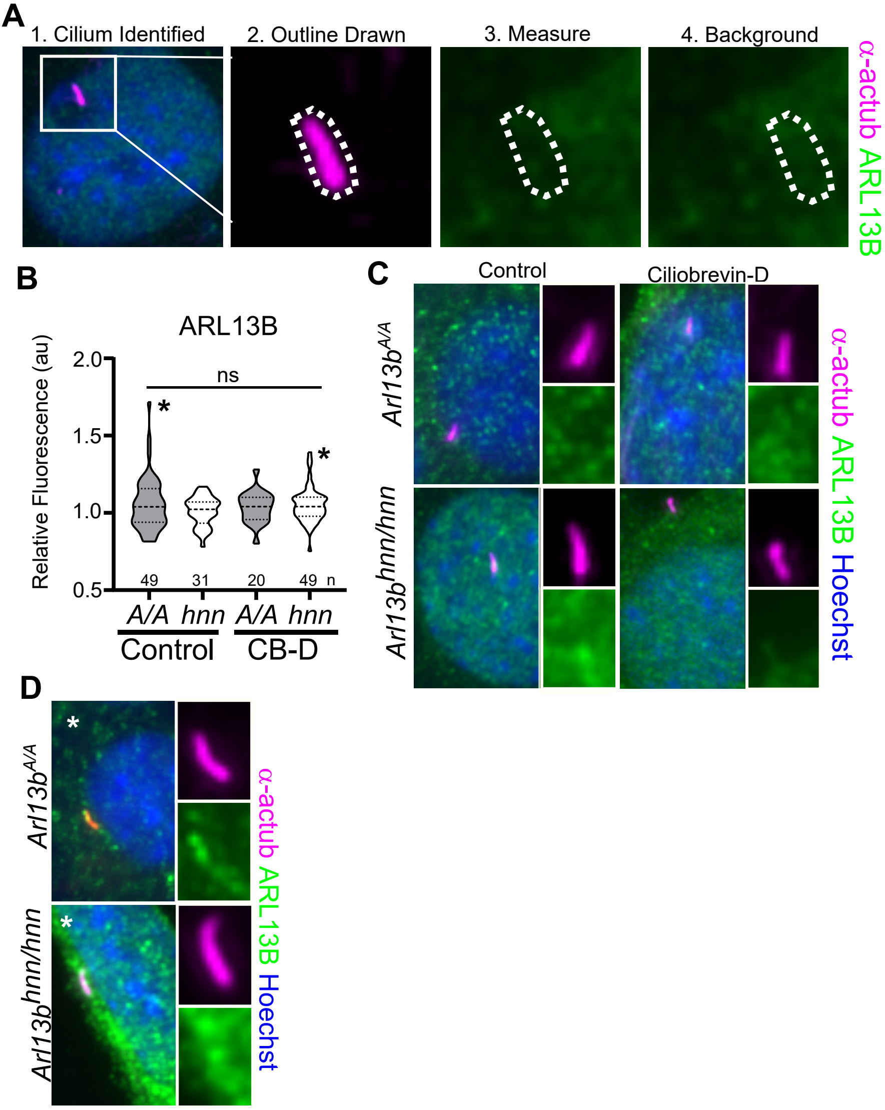

### Figure 3 Supplement 2

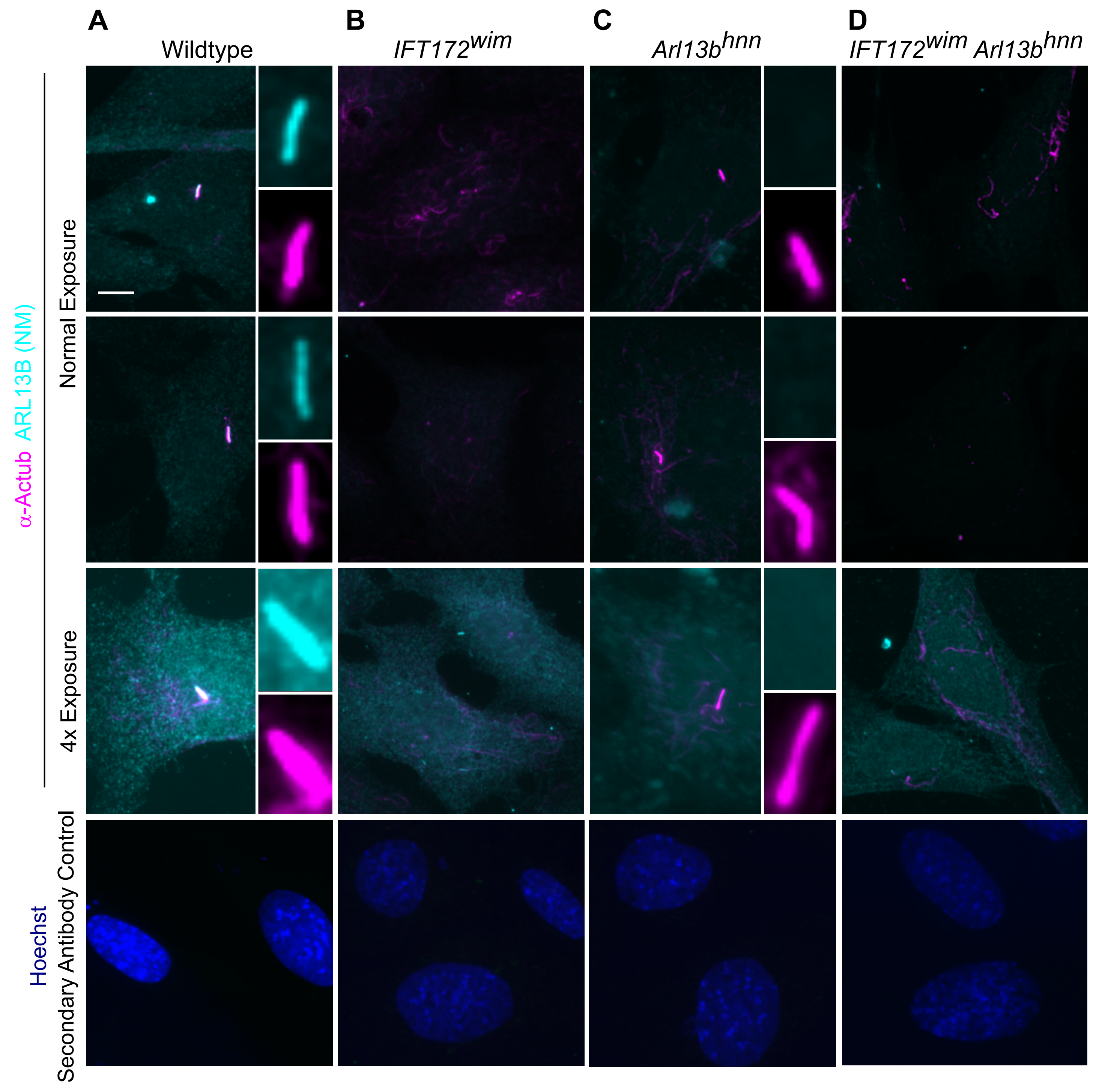
